# Pharmacological and Transcriptomic Exploration of β2-Adrenergic Receptor-Gα15 Signaling in THP-1-Derived Macrophages

**DOI:** 10.64898/2026.02.09.704723

**Authors:** Yuan-En Sun, Qing Li, Justin G English

## Abstract

Myocardial infarction and heart failure are leading global causes of mortality. Chronic β-adrenergic receptor (βAR) activation in cardiomyocytes promotes heart failure via Gαs signaling after myocardial infarction, whereas β2AR activation may also provide cardiac protection and repair through alternative pathways. Macrophages play a pivotal role in cardiac repair, and β2AR has been reported to signal via the hematopoietic-specific Gα15 in these cells. We aimed to characterize signaling bias between Gαs and Gα15 downstream of β2AR and to elucidate their roles in macrophage polarization. Using TRUPATH triple assays, we observed that several β2AR agonists activate Gα15 with at least an order of magnitude greater potency than Gαs in this system. In addition, clinically used β-blockers may exhibit differential inhibition on these two pathways. Macrophages are briefly classified into M1 and M2 polarization according to their activating stimuli and functional properties. Transcriptomic profiling of THP-1-derived macrophage-like cells treated with the β2AR agonist clenbuterol revealed enrichment of M1 transcriptional profile and repair-related hallmarks. Knockdown of Gαs showed M1 enrichment, whereas Gα15 knockdown was associated with negative M2 enrichment as well as M1 enrichment. Loss of either Gαs or Gα15 negatively affected repair-associated hallmarks. In contrast, pharmacological intervention of the Gαs-cAMP signaling produced opposing M1/M2 transcriptional responses, while suppressing repair-associated hallmarks. These *in vitro* findings explore the distinct pharmacological profiles of β2AR ligands toward Gαs and Gα15 and reveal how β2AR agonism may modulate macrophage function through dual-transducer signaling.

**Significance statement:** β2AR agonism engaged Gα15 signaling in the TRUPATH triple assay, with several agonists showing higher apparent potency for Gα15 than Gαs and β-blockers showing distinct inhibitory profiles across Gαs and Gα15. In macrophage-like cells, β2AR was associated with an M1 transcriptional profile. Notably, Gα15 may contribute to M2-associated transcriptional profile under β2AR activation, whereas the Gαs-cAMP showed an opposing M1 transcriptional pattern. These findings suggest that Gα15 signaling may represent a noncanonical pathway contributing to β2AR-induced transcriptional responses in macrophages.

## 1. Introduction

Cardiovascular diseases, such as myocardial infarction and congestive heart failure account for approximately 30% of annual deaths worldwide^1^. Among these, myocardial infarction, or heart attack, is the leading cause of heart disease-related mortality. Myocardial infarction is defined as myocardial necrosis resulting from sustained ischemia, mostly caused by acute obstruction of coronary blood flow^2^. Patients experiencing myocardial infarction lead to rapid ischemic tissue damage of heart muscle, loss of cardiomyocytes, reduced cardiac contractility^3^. To compensate, the sympathetic nervous system releases catecholamines to raise heart rate and promote vasoconstriction, but long-term elevation becomes cardiotoxic by driving oxidative stress, increasing oxygen demand, and promoting fibrosis, ultimately leading to heart failure^4, 5^. Although pharmacological intervention of myocardial infarction and heart failure with reduced ejection fraction has improved clinical outcomes, the development of heart failure following myocardial infarction remains high^6, 7^.

Catecholamines targeting β1 & β2-AR raise heart rate after myocardial infarction. However, sustained β1 & β2AR activation leads to apoptosis of cardiomyocytes, establishing a deleterious loop of heart damage and progressing to HF^8, 9, 10^. β-blockers, which antagonize β1 & β2AR-Gαs signaling by blocking the binding of catecholamines, are the standard treatment for patients after myocardial infarction to prevent the development of heart failure and have been used for over 20 years due to their ability to reduce mortality and prevent arrhythmias^11^. Among β-blockers, carvedilol has been reported as the most effective agent for reducing mortality following myocardial infarction^12^.

Although sustained β1 & β2AR-Gαs signaling in cardiomyocytes is deleterious, β2AR agonism has been demonstrated to benefit heart function and show cardioprotective effects^13, 14, 15^. Besides coupling to Gαs, it has been demonstrated that β2AR can switch coupling to Gαi/o and recruit β-arrestins to activate different signaling pathways^16, 17, 18^. Cardioprotective effects from β2AR via Gαi/o and β-arrestins have been proposed^19, 20, 21, 22^. Given the beneficial effects of alternative β2AR signaling pathways in cardiomyocytes, ongoing studies aim to define the underlying mechanisms that β2AR signaling contributes to cardioprotection and to evaluate its therapeutic potential. In addition to Gαi/o and β-arrestins, β2AR has been demonstrated to activate Gα15 by measuring its downstream signaling activity and the dissociation of Gα15 from β2AR using a BRET sensor^23, 24^. Gα15 is classified as a member of the Gαq/11 subtype and is known to induce downstream Ca2+ mobilization, IP3, and DAG production^25^. Gα15 is exclusively expressed in blood cells. The function of Gα15 has been proposed to be involved in cell proliferation and maturation^26, 27^. However, mice lacking Gα15 exhibit normal hematopoiesis and inflammatory responses, suggesting that the absence of Gα15 signaling is functionally compensated for by other Gαq and Gαi proteins^28^. Therefore, the exact function of Gα15 in blood cells remains poorly characterized.

The prevailing studies for cardioprotective β2AR signaling has centered on cardiomyocyte activity. However, many different cell types in the heart express β2AR, and it has been demonstrated that cell-type specific β2AR signaling can have a distinct contribution to cardiac function^29^. Immune cells have been demonstrated to play an important role in heart repair and recovery after myocardial infarction^30^. The early inflammatory phase occurs after myocardial infarction, resulting in the induction of pro-inflammatory immune cells to remove the damaged tissue, followed by the repair phase with the resolution of inflammation and repair of the heart. It is shown that the post-myocardial infarction inflammation remains unresolved, resulting in chronic inflammation of the heart, which causes sustained damage and eventually progresses to heart failure^31^. Among immune cells, macrophages have been extensively studied in the context of heart disease. Macrophages with anti-inflammatory phenotypes play a key role in resolving inflammation and promoting wound healing after myocardial infarction^32, 33^. Depletion of anti-inflammatory macrophages has been shown to impair cardiac function post myocardial infarction^34^. Conversely, their presence contributes to improved cardiac function and repair^35, 36^. Macrophages activity can be regulated by GPCR-G protein signaling. Gαs mainly signals through adenylyl cyclase-cAMP pathways and has been implicated in macrophage immune responses, foam-cell formation, and inflammasome regulation, whereas Gα15 induces PLC/ Ca^2+^ signaling and contributes to inflammatory receptor responses such as C5a-mediated Ca^2+^ mobilization in macrophage. Together, these pathways may determine how macrophages respond via GPCR signaling and provide opportunities to selectively modulate macrophage function through biased signaling^37, 38, 39, 40, 41^. Furthermore, β2AR in the immune cells has been reported to be essential for survival after myocardial infarction, which suggests that β2AR signaling in macrophages might contribute to cardiac repair^41^.

β2AR signaling in macrophages has been reported to exert context-dependent immunomodulatory effects^42^, characterized by increased *IL1b* and *IL6* expression, attenuated TNF-α secretion, and enhanced IL-10 production^43, 44, 45^. However, whether the macrophage activities induced by β2AR agonism are associated with cardioprotection is still not fully studied. In macrophages, β2AR may couple to the Gα15 signaling pathway, potentially giving rise to cellular responses that differ from canonical Gαs-mediated signaling. In addition, the pharmacological profiles of clinically prescribed β2 agonists and β-blockers on β2AR-mediated Gα15 activation remains entirely uncharacterized. Elucidating the pharmacology of this pathway and the impact on macrophages may provide valuable insights for the potential therapeutics on MI. Here, we conducted an exploratory study for the pharmacological profile of β2AR agonists and β-blockers on Gαs and Gα15 activation and associated macrophage transcriptional responses. In TRUPATH triple sensors, we observed that β2AR agonists activated Gα15 with higher pEC_50_ than Gαs, while clinically prescribed β-blockers showed differential blockade on β2AR-mediated Gαs and Gα15 signaling in the presence of an EC_80_ concentration of epinephrine. We further used human THP-1-derived macrophage-like cells as an in vitro model to examine transcriptional profile associated with β2AR activation as well as Gαs and Gα15 signaling. These analyses suggest that β2AR signaling through Gαs and Gα15 is associated with distinct macrophage-related transcriptional profiles; however, these findings should be interpreted as exploratory study rather than definitive evidence of functional macrophage polarization.

## 2. Materials and Methods

### 2.1 Plasmids

NTSR1 and β2AR were cloned into pcDNA3.1(+) (V79020, Invitrogen). Gαs and Gα15 TRUPATH Triple sensors used in this study were from Addgene (https://www.addgene.org/browse/article/28233562/). The TRUPATH triple Gαs-short construct was created via PCR amplification of the Gαs-RLuc and Gβ-IRES-GFP-Gγ components of the TRUPATH Triple plasmid, and restriction enzyme digest-mediated subcloning to the pCMV-BI plasmid (Addgene #126475). Lentiviral plasmids: pCMV-VSV-G, psPax2, and pLKO-mEmerald used in this study were from Dr. Roh-Johnson’s lab at the University of Utah. RFP shRNA: 5’ CTCAGTTCCAGTACGGCTCCA 3’ (Plasmid #125778, Addgene)^46^, Gαs shRNA: 5’ TTGCTTTGTTAATCATGCCCTA 3’, and Gα15 shRNA: 5’ TTTTGAACCAGGGTAGTTCCAG 3’^47^ were cloned into pLKO-mEmerald. pLVX-GαsBP2-mKate2 of wild-type and mutants were from Dr. Garcia-Marcos’ lab. All constructs were verified by Sanger sequencing at the University of Utah core facility and Nanopore sequencing with Plasmidsaurus.

### 2.2 Ligands

Neurotensin 8-13 (Cat# 24718, Cayman Chemical), (−)-epinephrine (+)-bitartrate salt (Cat#324900, Sigma), formoterol hemifumarate (Cat#1448, Tocris), fenoterol hydrobromide (Cat#S5768, Selleckchem), clenbuterol hydrochloride (Cat#C5423, Sigma), terbutaline hemisulfate salt (Cat#T2528, Sigma), carvedilol (Cat#2685, Tocris), labetalol Hydrochloride (Cat#PHR1335, Sigma), and timolol Maleate (Cat#PHR2593, Sigma) were dissolved in DMSO to prepare 10 mM stock solutions and stored at −80 °C until use.

### 2.3 Cell culture

HEK293 cells (ATCC# CRL-1573) were cultured in Dulbecco’s Modified Eagle Medium (DMEM) (cat #10566024, ThermoScientific), supplemented with 10% fetal bovine serum (FBS) (Cat # FB-11, Omega Scientific) or 1% dialyzed FBS (Cat # FB-03, Omega Scientific), penicillin-streptomycin (0.1mg/mL) (Cat # 15140163, ThermoScientific) at 37°C with 5% CO2. THP-1 cells (from Dr. Roh-Johnson’s lab at the University of Utah) a human monocytic leukemia cell line were cultured in RPMI 1640 medium (Cat #11875119, ThermoScientific), supplemented with 10% FBS, 5mL 2M HEPES, 500µL 55mM β-mercaptoethanol, and 2.5mL penicillin-streptomycin at 37°C with 5% CO2.

### 2.4 TRUPATH Triples BRET2 Assay

HEK293 cells were seeded at a density of 2×10^6 in 10mL DMEM in 10-cm plates overnight. Cells were transiently transfected with 1480ng of TRUPAHT triple sensor and 296ng receptor (5:1 ratio) using TransIT-2020 (Cat#MIR 5406, Mirus bio) and incubated overnight. After incubation, cells were harvested and resuspended in 1% dialyzed FBS DMEM. For agonists, cells were then plated at a density of 10,000 cells/40μL for Gα15 and 20,000 cells/40μL for Gαs per well in the PDK (Sigma, P6407-5MG)-coated, clear-bottom 384-well assay plates (Greiner Bio-One, 781098) overnight. For antagonists, cells were plated at a density of 20,000 cells/40μL for both Gαs and Gα15 per well in 384-well assay plates overnight. Assay buffer (20 mM HEPES in 1x HBSS, pH to 7.4 with KOH) and 3x drug buffer (assay buffer + 3mg/ml BSA, 0.3 mg/ml ascorbic acid) were prepared at least 2 hours before the experiment. For agonists, drugs were diluted to 3x working concentration in 3x drug buffer using 96-well plates (Fisher, 12-566-120). Drug concentrations were created by serial dilution at half-log or log concentrations. For antagonists, epinephrine was prepared in 3x drug buffer at a 6x concentration corresponding to the EC_80_ for both Gαs and Gα15. Antagonists were prepared at a 6x working concentration in 3x drug buffer, and a series of antagonist concentrations was generated by log-scale serial dilution. Antagonists were then mixed with epinephrine in 96-well plates. The bioluminescent substrate Prolume purple (cat #369, NanoLight) was prepared in nanofuel (cat #399, NanoLight) to a concentration of 1 mM and stored at −80 °C. The growth media was flicked from each well, and a white backing (PerkinElmer, 6005199) was applied to the plate bottom. A concentration of 7.5 uM of the bioluminescent substrates, Prolume purple, was diluted in assay buffer, and each well of the plates was added 20uL of the substrate. Subsequently, 10μL of the drug solution was added to the well, producing a final well concentration of 5μM. The plate could be read by the PheraStar Reader. The optic settings on the PheraStar Reader were set to have one multichromatic and simultaneous dual emission. The well scan was set to have a 3mm spiral average. The top optic was selected in the Optic window. The setting time was 0.1s in the general settings. In the kinetic window, four cycles were read; the start time was 0s, and the interval time was 0.5s. All the wells were selected to be read, no matter how many were occupied. One cycle to read the entire plate was 335s. The time to normalize results was set to 0, and the pause before the next cycle was 1 second. In the measurement panel, the gain settings were set to 3800 for Gain A and 3000 for Gain B channels. Gain adjustment has a value of 40% for both the A and B channels. Cycle two or three data were used for analysis, and concentration–response curves were fit to a three-parameter logistic equation using GraphPad Prism.

### 2.5 THP-1 differentiation

THP-1 cells were seeded at a density of 2×10^6 in 2mL medium in 6-well plates and differentiated with 200ng/mL phorbol 12-myristate 13-acetate (PMA) (Cat#P1585, Sigma) for 72 hours. After 72 hours, the medium was replaced with PMA-free medium, and the cells were allowed to rest for 24 hours.

### 2.6 Immunostaining & Flow cytometry

Differentiated THP-1 cells were trypsinized with 1mL trypsin for 10 minutes and neutralized with 2mL PBS. Spin down at 300g for 5 minutes, and the supernatant was removed. Cells were resuspended in 100μL cell staining buffer (1x PBS + 2.5% BSA). 2.5μL of Human TruStain FcX™ Fc blocking solution (Cat#422301, Biolegend) was added to the cells. 1.25µL APC anti-mouse/human CD11b antibody (Cat#101211, Biolegend) was diluted in 5µL cell staining buffer. CD11b antibody was added to the cells and incubated at 4°C in the dark for 30 minutes. Cells were washed with 1mL washing buffer (1x PBS + 2% FBS + 0.1% Sodium Azide) twice, and resuspended in 300μL cell staining buffer. Cells were then transferred for testing by an Aria cell sorter. Flow-Jo V10 software was used for data analysis.

### 2.7 Lentiviral production & transduction

Viral packaging and transduction were modified from Greiner D. *et al*^48^. HEK293T cells were seeded at a density of 7×10^6 in 2mL medium in6 in 25mL DMEM in 15-cm plates. For SIV-VLPs, cells were transiently transfected with 1.69μg of pCMV-VSV-G and 9.57μg of pSIV3+ using TransIT-2020. For lentivirus containing shRNA or GαsBP2, cells were transiently transfected with 1.69μg of pCMV-VSV-G, 4.22μg of psPax2, and 5.35μg of pLKO-shRNA-mEmerald or pLVX-GαsBP2-mKate2, respectively, using TransIT 2020. After 24 hours of incubation, the medium was replaced with 30mL fresh DMEM. Cells were harvested between 36 to 48 hours after replacing the medium. The medium was collected and filtered through a 0.45μm filter. The viral particle was concentrated by ultracentrifugation at 26,000 RPM for 2 h at 4°C. The supernatant was removed. The viral particle was resuspended in 1000μL of cold RPMI 1640 medium by vigorously pipetting up and down (at least 40-50 times). The viral particle was incubated on ice for 30 minutes, followed by incubation at 4°C overnight. After incubation, the viral particle was pipetted up and down again and stored as aliquots at −80°C.

Differentiated macrophage-like cells were first incubated with SIV-VLPs (30μL/1×10^6 in 2mL medium in6 cells) for 30 minutes. After incubation, cells were incubated with 100μL of lentivirus for 48 hours. After 48 hours, the medium was replaced with fresh medium.

### 2.8 Immunoblotting

Cells were washed once with cold 1mL PBS and incubated with 150μL ice-cold RIPA buffer with protease inhibitors (PMSF, HALT, and Benzonase) (Cat#78429, Thermo Fisher) (Cat#E1014, Millipore Sigma) for 10 minutes. Cell lysate was transferred to a low-binding protein eppendorf and incubated on ice for 15 minutes. Spin down at max speed at 4°C for 10 minutes. Supernatants were transferred to clean low-binding protein tubes. Protein concentration was measured by Pierce BCA protein assay kit (Cat#23227, ThermoScientific). 10µg of crude cell lysate was used for SDS-PAGE gel running. Cell lysate was added with 4x Laemmli Sample Buffer (#1610747, Bio-Rad) that was spiked with 50mM DTT and heated at 98°C for 5 minutes. Samples were loaded on a 12-well 4-15% gradient gel (Cat#4561085, Bio-Rad). The gel was run at 180V for 1hour. Protein was transferred at 100V for 1 hour. The membrane was washed with DI water once and incubated with 10mL of blocking solution (1x TBS with 5% non-fat milk powder) in a Li-COR box, with agitation for 1 hour at room temperature. After discarding the blocking solution, the membrane was incubated with 5mL anti-Gαs rabbit primary antibody (Cat#10150-2-AP, Proteintech) at a 1:2000 dilution or anti-Gα15 rabbit primary antibody (Cat#NBP2-16557, Novus Biologicals) at a 1:1000 dilution in 1x TBS-T with 5% non-fat milk powder at 4°C overnight with agitation. The primary antibody solution was removed, and the membrane was washed 3 times with 10mL TBS-T for 5 minutes at room temperature. After washing, the membrane was incubated with 10mL secondary antibody, anti-rabbit IgG (H+L) (DyLight 800 4X PEG Conjugate) (Cat#5151P, Cell Signaling Technology) at 1:10,000 dilution, in 1x TBS-T with 5% non-fat milk powder at room temperature for 1 hour with agitation. The membrane was washed 3 times with 10mL TBS-T for 5 minutes at room temperature and first imaged by LI-COR Odyssey M for Gαs or Gα15. The membrane was then incubated with anti-GAPDH mouse primary antibody (Cat#MAB374, Millipore Sigma) at a 1:10,000 dilution at room temperature for 2 hours as a reference for normalization, followed by the washing step and incubation with 10mL secondary antibody, anti-mouse IgG (H+L) (DyLight 680 Conjugate) (Cat#5470P, Cell Signaling Technology) at 1:10,000 dilution, in 1x TBS-T with 5% non-fat milk powder at room temperature for 1 hour with agitation. The membrane was washed 3 times with 10mL TBS-T for 5 minutes at room temperature and imaged by LI-COR Odyssey M for GAPDH. The protein expression level was quantified by ImageJ.

### 2.9 RNA sequencing

Cells were treated with 20μL of 100μM Clenbuterol (final concentration is 1μM) and vehicle for 3 hours at 37°C with 5% CO2. After 3 hours, Actinomycin D (Cat#A1410, Millipore Sigma) was added to the cells to a final concentration of 15μM to terminate transcription and incubated at 37°C with 5% CO2 for 10 minutes. Cells were washed with 1mL pre-warm PBS once and incubated with 1mL pre-warm trypsin for 10 minutes. Cells were neutralized with 2mL PBS and spun down at 300g for 5 minutes. The supernatant was aspirated, and cells were resuspended in 250μL cell sorting buffer with 15μM Actinomycin D. Cells were kept on ice and sorted by Aria cell sorter for GFP or RFP-positive cells. GFP & RFP-positive cells were collected in 500μL RNAlater solution (Cat#AM7020, ThermoScientific) on ice. RNA was extracted using the Zymo Direct-Zol Mini Kit (Cat#R2050, Zymo Research). RNA samples were stored at −80°C. Intact poly(A) RNA was purified from total RNA samples (10-500 ng) using the NEBNext Poly(A) mRNA Magnetic Isolation Module (E7490). The library was prepared using the NEBNext Ultra II Directional RNA Library Prep Kit for Illumina (E7760L). Purified libraries were qualified on an Agilent Technologies 4150 TapeStation using a D1000 ScreenTape assay (cat# 5067-5582 and 5067-5583). The molarity of adapter-modified molecules was defined by quantitative PCR using the Kapa Biosystems Kapa Library Quant Kit (cat#KK4824). Individual libraries were normalized to 5nM in preparation for Illumina sequence analysis. Sequencing libraries were chemically denatured in preparation for sequencing. Following transfer of the denatured samples to an Illumina NovaSeq X instrument, a 151 x 151 cycle paired-end sequence run was performed using a NovaSeq X Series 10B Reagent Kit (20085594).

### 2.10 Bioinformatics analysis

For analysis of differential expression, the human GRCh38 FASTA and GTF files were downloaded from Ensembl release 112, and the reference database was created using STAR version 2.7. 9a with splice junctions optimized for 100 base-pair reads (Dobin et al., 2013). Optical duplicates were removed from the paired-end FASTQ files using BBMap’s Clumpify utility (v38.34), and reads were trimmed of adapters using cutadapt 1.16. The trimmed reads were aligned to the reference database using STAR. Mapped reads were assigned to annotated genes in the GTF file using featureCounts version 1.6.3. The output files from cutadapt, FastQC, STAR, and featureCounts were summarized using MultiQC to check for sample outliers. Read counts were normalized and analyzed using the Bioconductor DESeq2 analysis package (v1.42.1). Differentially expressed genes (DEGs) were defined as genes with an adjusted P value < 0.05, without applying an additional log2 fold change cutoff. All DEGs from DESeq2 analysis were subjected to gene set enrichment analysis (GSEA) against the Hallmark gene sets and Immunologic signature gene sets from Molecular Signatures Database (MSigDB)^49, 50, 51^, and the top ten hallmarks were listed. For exploratory analyses, enrichment results meeting a false discovery rate (FDR) threshold of < 0.25 were considered significant. In addition, DEGs meeting an adjusted P value < 0.05 were used for gene ontology (GO) enrichment analysis, and the top five enriched GO terms were listed^52^. GSEA and GO enrichment analyses were conducted in R using the fgsea and clusterProfiler packages, respectively.

### 2.11 Statistics and analysis

All cell-based and biochemical experiments were performed with three independent biological replicates unless otherwise stated. For the TRUPATH triple-sensor assay, three independent biological experiments were conducted, each with four technical replicates. Concentration-response curves are shown as mean ± SEM from the indicated number of independent experiments and were fitted in GraphPad Prism 10 using nonlinear regression with a three-parameter variable-slope model, with the Hill slope constrained to −1. Pharmacological parameters (pEC_50_, pIC_50_, efficacy, and Inhibition) are presented as mean ± SD. Statistical analyses of pharmacological parameters were performed using values derived from three independent biological replicates. One-way ANOVA followed by Dunnett’s test was used when comparing multiple groups with a single control, Tukey’s test was used for all pairwise comparisons, and two-way ANOVA followed by Šídák’s test was used for selected planned comparisons involving two independent variables. P value < 0.05 was considered statistically significant. All experiments and analyses were conducted in an exploratory framework without a prespecified formal null hypothesis, statistical values should be interpreted as descriptive rather than support formal hypothesis testing, and the resulting findings should be considered hypothesis-generating pending independent validation.

## 3. Results

### 3.1 β2AR agonists are more potent activators of Gα15 than Gαs

Gα15 has been demonstrated to promiscuously couple to many GPCRs. β2AR has been reported to couple to Gα15 by measurement of downstream second messenger and dissociation from β2AR using TRUPATH BRET sensors^23, 24^. These suggest that Gα15 may serve as an alternative signaling pathway of β2AR in the cell expressing Gα15, such as immune cells. The pharmacological characterization of β2AR agonists and antagonists has priactically focused on canonical Gαs-mediated signaling pathways. To further investigate β2AR-Gα15 signaling, we first characterized the pharmacological profiles of few β2AR agonists activating Gα15 and compared them with Gαs. These β2AR agonists have been previously reported to exert cardioprotective effects or wound healing, including formoterol, fenoterol, clenbuterol, and terbutaline^53, 54, 55, 56^. We first confirmed epinephrine-induced β2AR activation on Gα15 using the TRUPATH triple sensors developed by our laboratory. TRUPATH is a BRET-based biosensor platform used to monitor G protein activation in live cells following ligand-stimulated GPCR activation. The β2AR-Gα15 response was benchmarked against NTSR1-Gα15, which served as a positive control for Gα15. β2AR activated Gα15 with a pEC_50_ of 9.004 ± 0.222, whereas NTSR1 produced a pEC_50_ of 10.066 ± 0.133. The maximal efficacy of β2AR-mediated Gα15 activation was approximately 85% of that observed for NTSR1 (Supplemental Figure 1A). Next, we characterized the pharmacological profiles of additional four β2AR agonists for Gαs and Gα15 using the TRUPATH triple sensors (Supplemental Figure 1B-F). The concentration-response curves of each agonist were normalized to epinephrine, which served as the reference agonist for both Gαs and Gα15 (Fig. 1A & B). The heat map depicts the pEC_50_ and relative efficacy of all agonists for Gαs and Gα15 activation (Fig. 1C). We observed that all agonists activated Gα15 with pEC_50_ values at least one log unit higher than their corresponding pEC_50_ values for Gαs activation, as shown in Fig. S1B-F. Among all agonists tested, terbutaline exhibited the lowest pEC_50_ values for both Gαs and Gα15 activation, consistent with a previous report on its binding affinity^57^ (Supplemental Figure 2A). In addition, terbutaline displayed about 50% reduced relative efficacy for Gαs activation, but not for Gα15 activation, compared with epinephrine (Supplemental Figure 2B). This finding suggests that, unlike the other agonists tested, terbutaline may act as a full agonist for β2AR-mediated Gα15 activation while functioning as a partial agonist for Gαs activation. Altogether, the β2AR agonists tested elicited concentration-dependent activation of both Gαs and Gα15 in TRUPATH triple sensor assay. Generally higher pEC_50_ values were observed for Gα15 activation than for Gαs activation under this experimental condition. However, since Gαs and Gα15 activation were assessed using distinct sensors, these apparent pEC_50_ differences should be interpreted cautiously and may reflect assay-dependent factors rather than intrinsic differences in G protein coupling.

**Figure 1.**
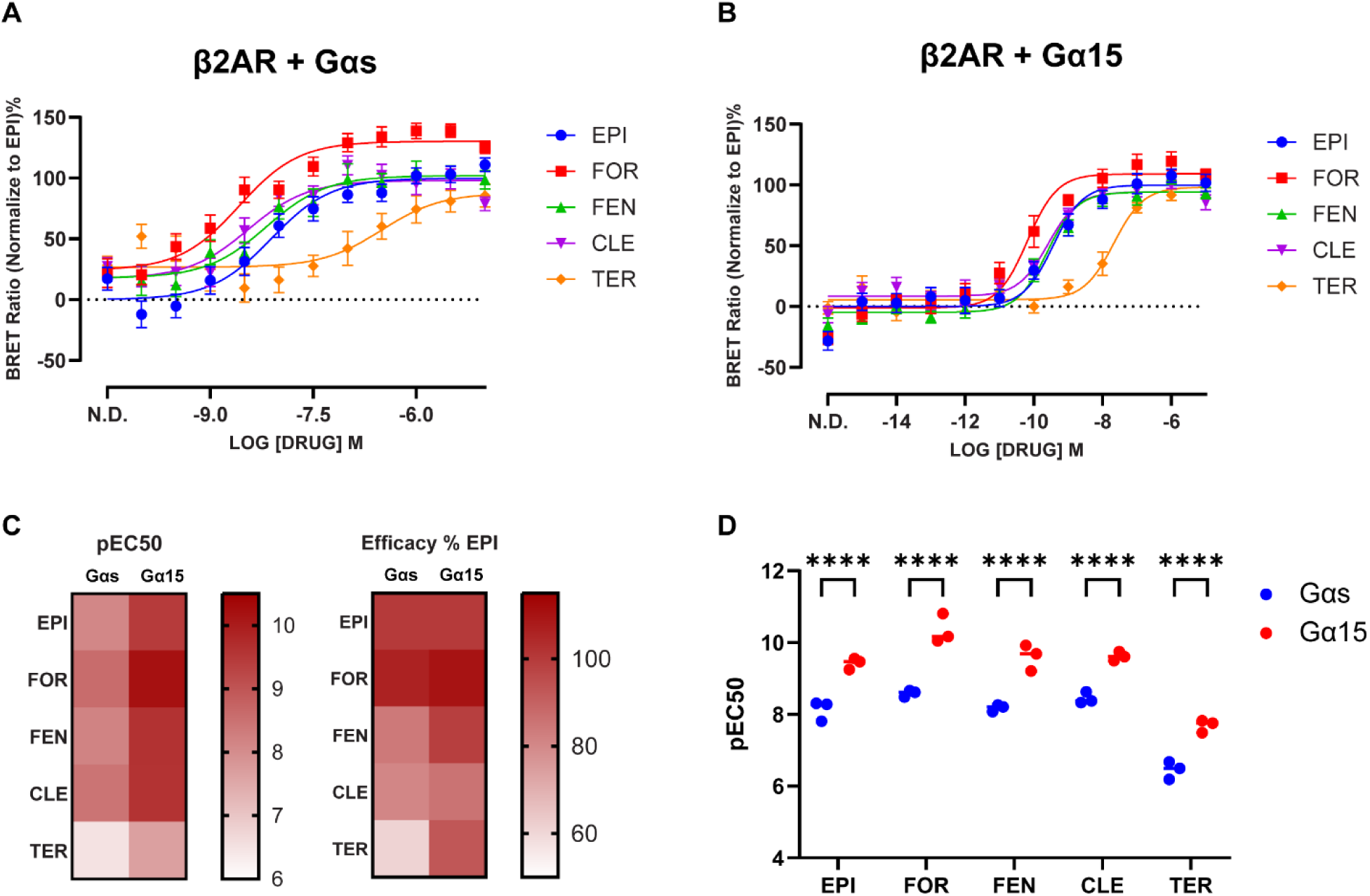
β2AR agonists are more potent activators of Gα15 than Gαs. (A) Gαs TRUPATH concentration-response curves of β2AR with epinephrine (EPI), formoterol (FOR), fenoterol (FEN), clenbuterol (CLE), and terbutaline (TER). Data are represented as mean values ± SEM, N=3, n=4. (B) Gα15 TRUPATH concentration-response curves of β2AR with epinephrine, formoterol, fenoterol, clenbuterol, and terbutaline. Data are represented as mean values ± SEM, N=3, n=4. (C) Heat map of the potency (pEC50) and relative amplitude (efficacy % EPI) of β2AR agonists measured in panels A & B, for both GαS and Gα15. Heatmap values represent mean values. (D) Statistical analysis of pEC_50_ Values for Gαs and Gα15 across all drugs. Two-way ANOVA followed by Šídák’s test, P-value: 0.1234(ns), 0.0332(*), 0.0021(**), 0.0002(***) & 0.0001(****). N refers to biological replicate, and n refers to technical replicate. pEC_50_ and efficacy values are presented as mean ± SD from three independent biological replicates.

### 3.2 β2AR antagonists exhibit differential Inhibition of Gαs and Gα15

Several β-blockers are used to prevent heart failure after myocardial infarction due to their ability to block β1 and β2AR-Gαs signaling in cardiomyocytes. However, the inhibitory potency of β-blockers toward β2AR-Gα15 signaling remains unclear, and differences in potency may result in differential modulation of β2AR signaling in immune cells. To investigate this, we selected three non-selective β-blockers that lack β-agonist activity^58^—carvedilol, labetalol, and timolol—based on their distinct clinical indications. Carvedilol and labetalol are also known to exhibit additional blockade of α1-adrenergic receptor signaling, whereas timolol only blocks β1 and β2Ars^59^. Clinically, Carvedilol is mainly used for myocardial infarction and heart failure, labetalol is used for treating hypertension, and timolol is used for glaucoma^12, 59, 60^. However, they have all been demonstrated to be beneficial in heart function after myocardial infarction in mouse model^12, 61, 62^. We characterized the pharmacological profiles of the three β-blockers by assessing their inhibitory effects on epinephrine-induced Gαs and Gα15 activation using the TRUPATH triple assay. For each G protein, epinephrine was applied at the previously established EC80 concentration (Supplemental Figure 1B). BRET ratio values at each concentration of all β-blockers were normalized to the vehicle (N.D.) for both Gαs and Gα15. (Fig. 2A & B). Under epinephrine stimulation at EC_80_, we observed carvedilol pIC_50_: 7.318 ± 0.216, labetalol pIC_50_: 7.025 ± 0.167, and timolol pIC_50_: 8.567 ± 0.428 for Gαs; carvedilol pIC_50_: 7.151 ± 0.165, labetalol pIC_50_: 7.128 ± 0.012, and timolol pIC_50_: 7.787 ± 0.038 for Gα15. Carvedilol and labetalol exhibited no difference in pIC_50_ between Gαs and Gα15, whereas timolol displayed a higher pIC_50_ for Gαs compared with Gα15 (Fig. 2C). pIC_50_ of timolol for both Gαs and Gα15 are higher than carvedilol and labetalol (Fig. 2D). Furthermore, labetalol showed significantly lower inhibitory efficacy for the Gα15 activation than carvedilol and timolol. Labetalol showed similar means of inhibitory efficacy to carvedilol and timolol for Gαs. However, the replicates of carvedilol and timolol were highly variable, these differences did not reach statistical significance, and the data do not support a definitive distinction between labetalol and the other two β-blockers for inhibition of Gαs in this experiment (Fig. 2E). Together, we observed that the β-blockers tested may exhibit distinguishable inhibitory profiles toward β2AR-mediated Gαs and Gα15 activation under epinephrine-stimulated conditions. However, since these findings were obtained only using the TRUPATH triple assay, they should be interpreted with caution and will require further validation in other assays.

**Figure 2.**
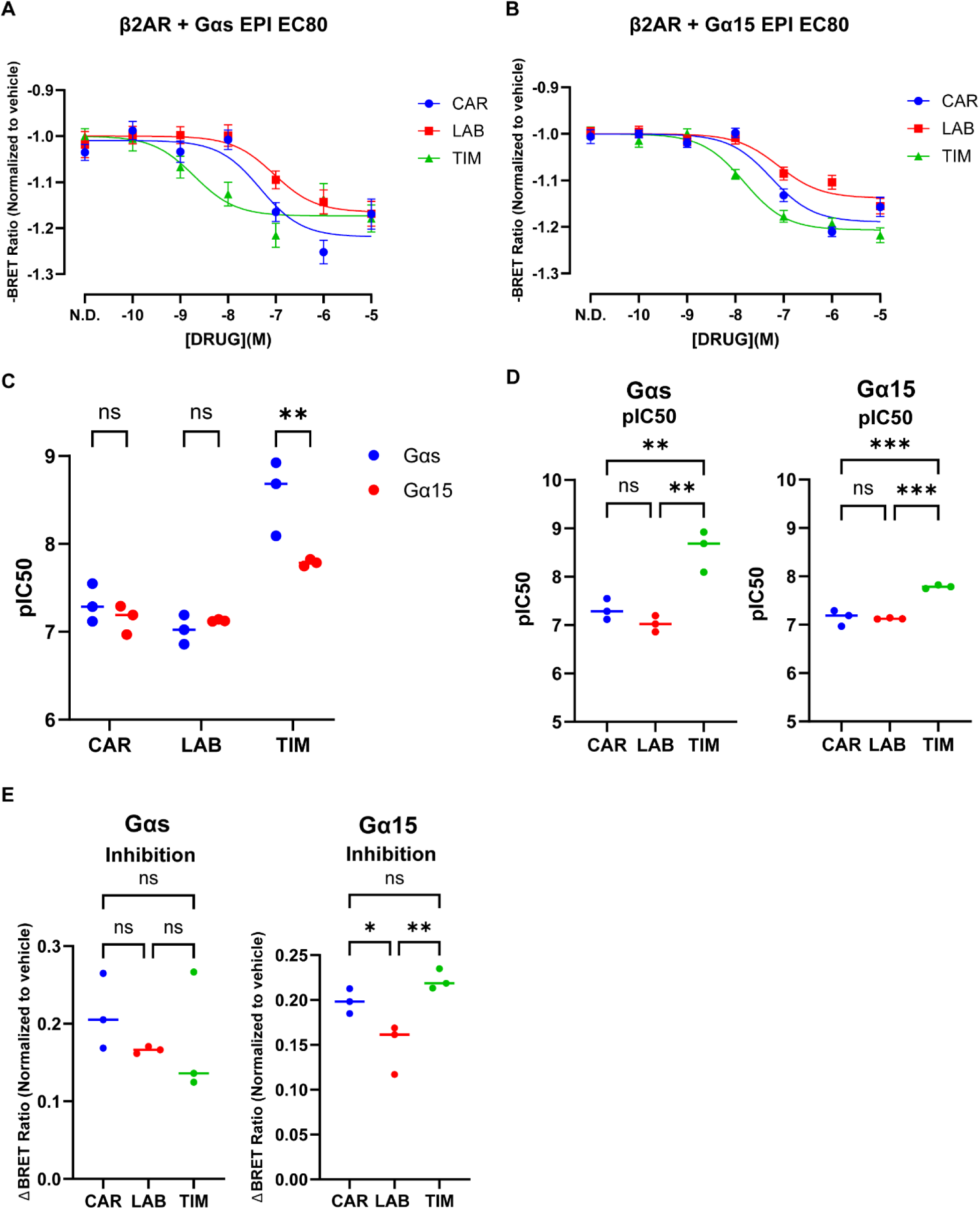
Inhibition of Gαs and Gα15 by β2AR antagonists at EC_80_ of epinephrine. (A) Gαs TRUPATH concentration-response curves of β2AR with carvedilol (CAR), labetalol (LAB), and timolol (TIM) at EC_80_ of epinephrine of Gαs. Data are represented as mean values ± SEM, N=3, n=4. (B) Gα15 TRUPATH concentration-response curves of β2AR with carvedilol, labetalol, and timolol at EC_80_ of epinephrine of Gα15. Data are represented as mean values ± SEM, N=3, n=4. (C) Statistical analysis of the potency of inhibition (pIC_50_) in panels A & B between Gαs and Gα15 across all antagonists. Two-way ANOVA followed by Šídák’s test. (D) Statistical analysis of pIC_50_ between antagonists for both Gαs and Gα15. One-way ANOVA followed by Tukey’s test. (E) Statistical analysis of the relative inhibition between antagonists for both Gαs and Gα15. One-way ANOVA followed by Tukey’s test. P-value: 0.1234(ns), 0.0332(*), 0.0021(**) & 0.0002(***). N refers to biological replicate, and n refers to technical replicate. pIC_50_ and inhibition values are presented as mean ± SD from three independent biological replicates.

### 3.3 Knockdown of Gαs and Gα15 in THP-1-derived macrophage-like cells

The TRUPATH triple assay revealed that β2AR agonists activated Gα15 signaling, whereas β-blockers effectively attenuated epinephrine-induced Gα15 activation. Consistently, a previous study reported that the β2AR agonist, isoprenaline, stimulates inositol phosphate accumulation in THP-1 cells^23^, and other studies have also demonstrated that β2AR activation promotes PLC activity in erythrocytes^63, 64^, thereby providing evidence for endogenous coupling between β2AR and Gα15 in the cell natively expressing Gα15. Nevertheless, the precise physiological role of β2AR-Gα15 signaling in hematopoietic cells remains unclear, as does the extent to which cellular responses differ from β2AR-Gαs versus β2AR-Gα15 signaling.

According to single-cell transcriptome analysis of the heart, Gα15 is mainly expressed in myeloid cells in the heart, including monocytes, macrophages, dendritic cells, and neutrophils^65^. Among these cell types, macrophages are the most studied cell type in heart diseases. Macrophages play an important role in mediating the inflammation after MI. Particularly, Anti-inflammatory macrophages resolve the inflammation and change the extracellular matrix to benefit heart function. β2AR agonism has been shown to mediate the inflammatory activities of macrophages. We hypothesized that β2AR agonism in macrophages may lead to signaling pathways associated with cardioprotective effects. To study β2AR signaling in macrophages, we used human THP-1 as the model, which is commonly used for macrophage studies^66^. THP-1 is leukemia monocytes and can be differentiated into a macrophage-like lineage. Following previous studies and testing, we established our own protocol to differentiate THP-1 cells for 72 hours with 200 ng/mL PMA, followed by a 24-hour rest period (Supplemental Figure 3A). After PMA treatment, the morphology of the cells changed dramatically, from suspension to adherent. Cells also expressed high level of CD11b, a macrophage marker^66^. In untreated THP-1 cells, ∼12% cells were CD11b-positive under basal conditions. PMA treatment markedly increased the proportion of CD11b-positive cells to ∼90%, indicating increased CD11b-positive cell population and supporting differentiation toward a macrophage-like cells. CD11b positivity was defined by flow cytometric gating relative to unstained samples (Supplemental Figure 3B). This result indicates that THP-1 can be successfully differentiated into macrophages with our protocol for further β2AR signaling studies in macrophage-like cells.

In macrophages, β2AR activates not only Gαs but also Gα15 signaling. Gα15 belongs to the Gαq/11 family and has been demonstrated to induce the Gαq/11 downstream signaling pathway, such as PLC activation, calcium mobilization, and IP3 and DAG production. Thus, Gαs and Gα15 signaling should have distinct signaling and transcriptional profile. To study how of β2AR signaling affect macrophage activity and the difference between Gαs and Gα15 signaling pathway in macrophages, we used short hairpin RNAs (shRNAs) to knock down Gαs and Gα15 into THP-1-derived macrophage-like cells by lentiviral transduction and performed RNA sequencing to demonstrate the transcriptional profile of β2AR as well as Gαs and Gα15 signaling under β2AR stimulation. Macrophage-like cells were transduced with shRNAs targeting either Gαs or Gα15 by lentiviral transduction. shRNA targeting red fluorescent protein (RFP) was used as the control. To evaluate knockdown efficiency and specificity, whole-cell lysates were analyzed by immunoblotting using antibodies against Gαs and Gα15. Gαs and Gα15 protein abundance was normalized to GAPDH from the same lane. GAPDH abundance did not differ significantly among experimental conditions (Supplemental Figure 3D). Quantification of the immunoblots showed that cells transduced with Gαs shRNA exhibited an approximately 80% reduction in Gαs protein abundance relative to the RFP shRNA control, with no statistically significant change in Gα15 expression. Conversely, cells transduced with Gα15 shRNA showed an approximately 40% reduction in Gα15 protein abundance relative to the RFP shRNA control, without significantly altering Gαs expression (Supplemental Figure 3C & E). These results indicate that the Gαs and Gα15 shRNAs used in this study selectively reduced the expression of their intended Gα protein targets.

### 3.4 Transcriptome analysis of β2AR signaling in macrophage-like cells

β2AR agonism in macrophages has been reported to stimulate pro-inflammatory and anti-inflammatory activities, though these observations are largely limited to analyses of cytokine secretions. However, the overall macrophage activity mediated by β2AR signaling in macrophages have not been well-studied yet. We aimed to use transcriptome analysis to study the transcriptional response induced by β2AR signaling in macrophage-like cells. Furthermore, we also investigated the downstream difference of transcriptional profile between Gαs and Gα15 signaling under β2AR stimulation by knocking down Gαs and Gα15 in cells. In parallel, we also exogenously expressed the newly developed Gαs peptide inhibitor, called GαsBP2, which blocks the interaction between Gαs and adenylyl cyclases, thereby preventing cAMP production, including in immune-cell contexts.^67^, as an orthogonal approach to block canonical Gαs-cAMP signaling. We selected clenbuterol, which is a more β2AR-selective agonist, to stimulate β2AR signaling in cells^57^.

Since the lentivirus encodes green and red fluorescent protein to indicate the presence of shRNA and the Gαs inhibitor, respectively, cells were sorted for GFP- or RFP-positive populations before RNA sequencing (Fig. 3A). Differential expression analysis (DEA) was performed to identify DEGs mediated by β2AR agonism in the clenbuterol-treated versus untreated groups in the presence of RFP shRNA and mKate2, which served as the respective controls for shRNA-mediated knockdown and Gαs inhibition (Supplemental Table 1 & 2). To identify the genes mediated by β2AR signaling confidently, we compared the DEGs of the RFP shRNA and mKate2 datasets with and without clenbuterol treatment and performed the correlation analysis (Fig. 3B & C). These results suggest that the DEGs identified in the two groups are strongly correlated and are primarily associated with clenbuterol-stimulated β2AR signaling. We observed that clenbuterol treatment significantly upregulated *NR4A1*, *NR4A3*, *NTRK1*, *NPTX1*, *SNAP25*, and *ADPRHL1*, while *CX3CR1* and *FFAR2* were significantly downregulated (Fig. 3C). Among these upregulated genes, *NR4A1*, *NR4A3*, and *NTRK1* have been reported to be involved in immune responses and regulation of inflammation^68, 69, 70^. In contrast, the downregulated gene, *CX3CR1* has also been implicated in macrophage recruitment^71^. Besides, we also observed upregulated expression of *IL10* and *IL1b*—classic anti- and pro-inflammatory cytokines, respectively—which have previously been reported to be mediated by β2AR signaling^72, 73^. Next, we performed GSEA on clenbuterol-induced genes identified in the RFP shRNA and mKate2 control conditions, together with GO term enrichment analysis of the shared genes in these two conditions. Both datasets showed enrichment of predefined MSigDB hallmark gene sets, including TNF signaling via NF-κB, inflammatory response, and IL-2/STAT5 signaling in GSEA. GO term enrichment analysis identified enrichment of terms associated with chemotaxis and small GTPase-mediated signal transduction (Fig. 3D). Together, these exploratory analyses suggest that clenbuterol-stimulated β2AR agonism is associated with immune-related transcriptional responses in THP-1-derived macrophage-like cells, for example, inflammatory signaling, immune regulatory, cytoskeletal remodeling, and migration.

**Figure 3.**
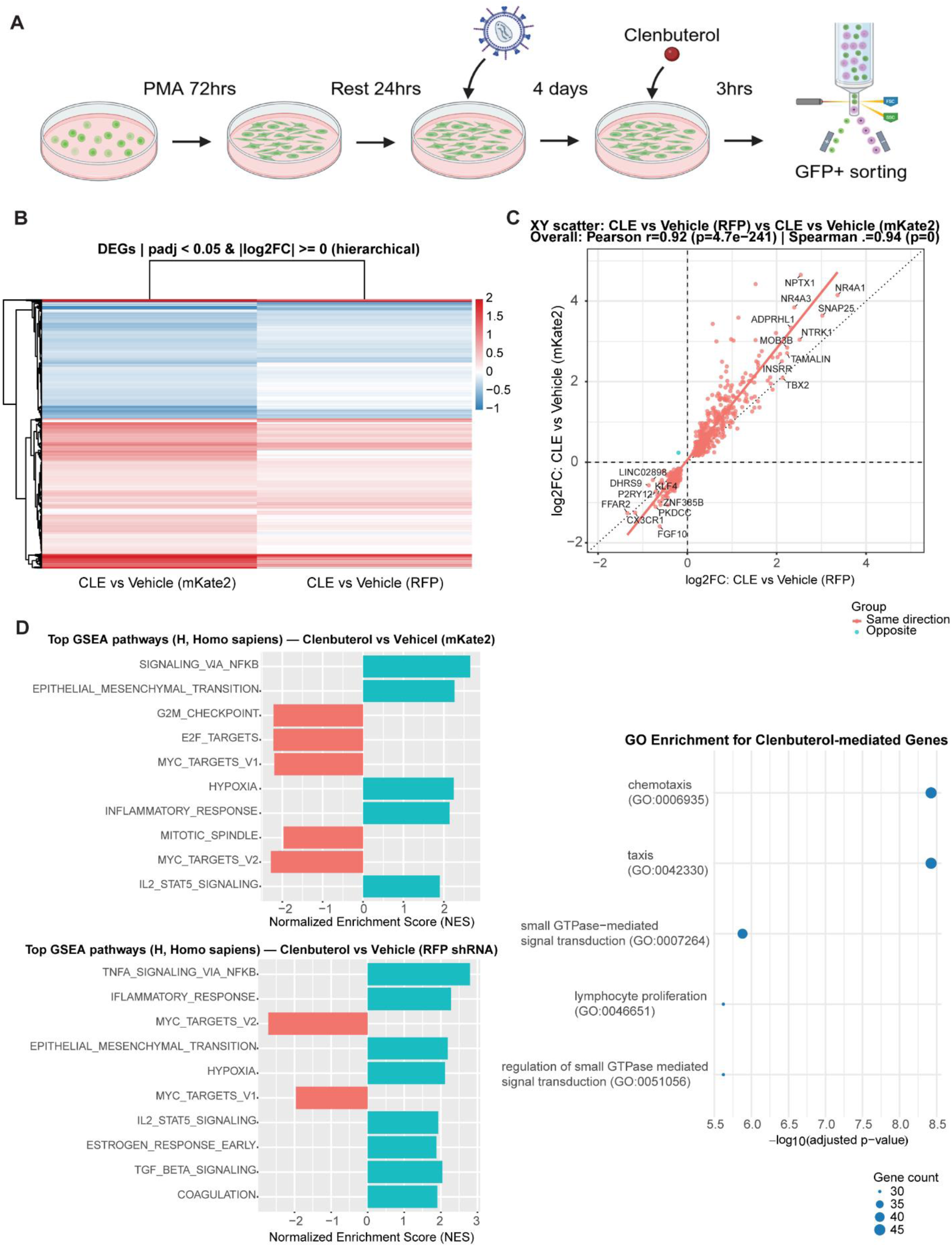
Transcriptomic profiling of clenbuterol-treated THP-1-derived macrophage-like cells. (A) The workflow of RNA sequencing for clenbuterol-treated THP-1-derived macrophage-like cells with Gαs and Gα15 knockdown. (B) Hierarchical clustering heatmap on DEGs from both RFP control groups and mKate2 groups, with and without clenbuterol treatment, defined by ANOVA with an adjusted p-value less than 0.05. N=3. (C) The correlation analysis of DEGs with adjusted p-values <0.05 from both RFP shRNA groups and mKate2 groups, with and without clenbuterol treatment. (D) The GSEA analysis of the clenbuterol-mediated genes in the context of expressing RFP shRNA and mKate2. (E) The GO term enrichment analysis of the clenbuterol-mediated genes. N refers to biological replicate.

### 3.5 Transcriptome analysis of β2AR-Gαs and Gα15 signaling in macrophage-like cells

To identify the downstream transcriptional profile of Gαs signaling under β2AR stimulation in macrophages, we performed the DEA for the Gαs knockdown group versus the RFP control group under clenbuterol treatment (Supplemental Table 3). We confirmed that *GNAS* expression was significantly reduced compared with control group (Fig. 4A). Canonically, β2AR activation couples to Gαs, which stimulates adenylyl cyclase, increases cAMP production, and induces downstream cAMP-responsive genes. Consistent with this pathway, the RNA abundance of previously reported β2AR-Gαs-cAMP/CREB responsive genes, including *NR4A1*, *NR4A2*, *NR4A3*, *FOS*, *DUSP1*, and *CREM*^74^ were reduced in Gαs knockdown groups compared with control groups under clenbuterol-treated conditions (Fig. 4A), supporting disruption of the β2AR-Gαs-cAMP/CREB signaling pathway by Gαs knockdown in the RNA-seq. We also performed the DEA for the GαsBP2 wild-type group versus the GαsBP2 mutant control group under clenbuterol-treated conditions (Supplemental Table 4). The DEA showed a similar reduction in these β2AR-Gαs-cAMP/CREB-responsive genes (Supplemental Figure 4A), supporting GαsBP2-mediated perturbation of the Gαs-cAMP signaling axis in the RNA-seq. However, since cAMP production and downstream signaling were not directly measured, these findings should be interpreted as transcriptional evidence consistent with perturbation of the Gαs-cAMP pathway, rather than direct biochemical evidence of cAMP inhibition in this experiment. We further compared the transcriptional profiles between Gαs knockdown and GαsBP2. Hierarchical clustering showed limited overlap between the Gαs knockdown and GαsBP2 transcriptional profiles (Supplemental Figure 4B). Together, these RNA-seq data of both Gαs knockdown and GαsBP2 can delineate downstream genes regulated by Gαs signaling. Notably, Gαs knockdown appears to exert broader effects on downstream transcription than specific blockade of Gαs-cAMP axis by GαsBP2.

**Figure 4.**
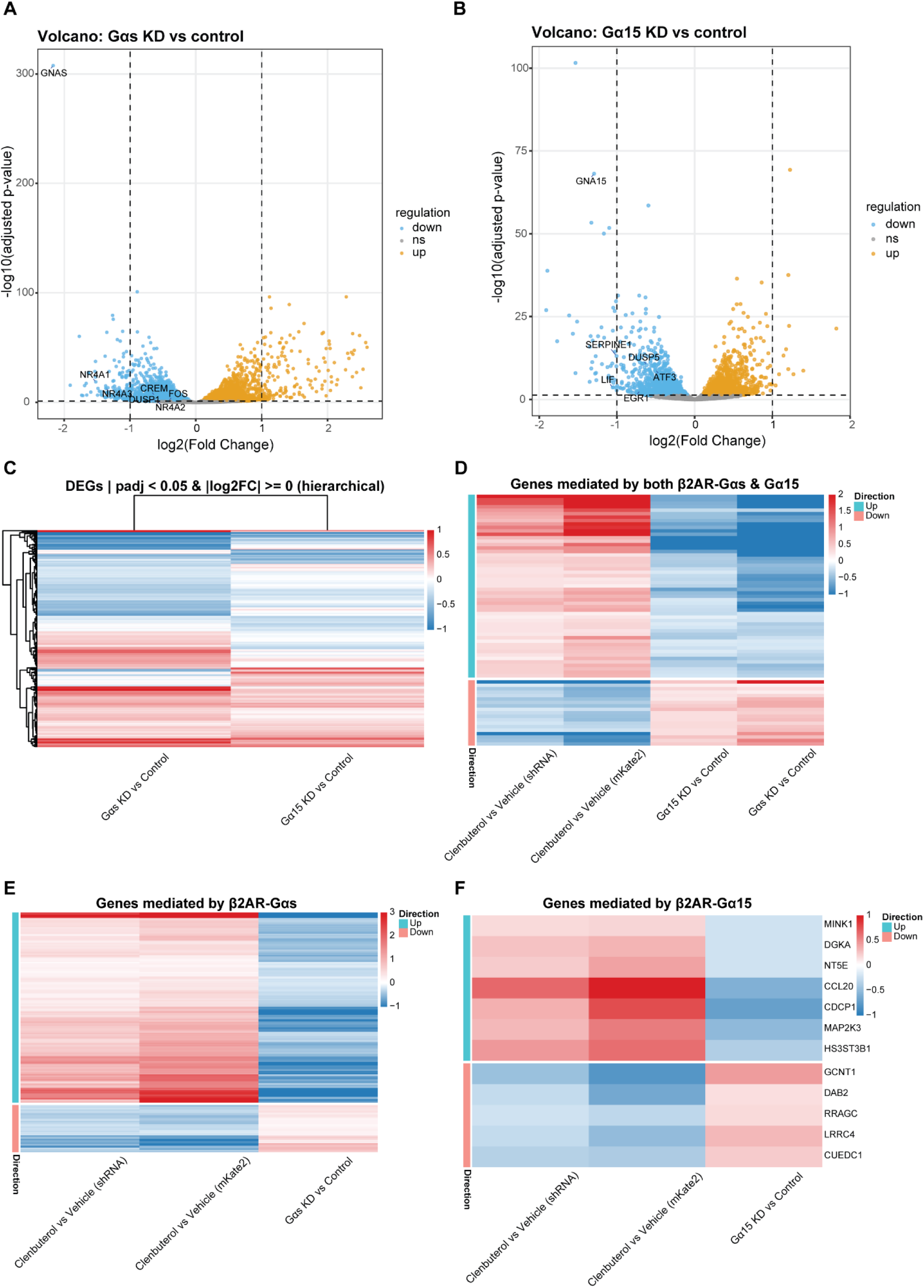
Transcriptomic Profiling of Gαs and Gα15-dependent signaling in response to clenbuterol in THP-1-derived macrophage-like cells. (A) Volcano plot of DEGs between the Gαs knockdown group versus control under clenbuterol treatment, with genes known to be regulated by the β2AR-Gαs-CREB pathway annotated, defined by ANOVA with an adjusted p-value <0.05 (yellow and blue dots). N=3. (B) Volcano plot of DEGs between the Gα15 knockdown group versus control under clenbuterol treatment, with genes known to be regulated by the Gαq/11 pathway annotated, defined by ANOVA with an adjusted p-value <0.05 (yellow and blue dots). N=3. (C) Hierarchical clustering heatmap on DEGs of Gαs or Gα15 knockdown versus control under clenbuterol treatment, defined by ANOVA with an adjusted p-value <0.05. (D) Heatmap of genes regulated by both β2AR-Gαs and −Gα15 signaling pathway. (E) Heatmap of genes uniquely regulated by β2AR-Gαs signaling pathway. (F) Heatmap of genes uniquely regulated by β2AR-Gα15 signaling pathway. N refers to biological replicate.

Subsequently, we want to characterize the downstream genes mediated by Gα15 signaling under β2AR stimulation in macrophages. We performed the DEA for the Gα15 knockdown group versus the RFP control group under clenbuterol treatment (Supplemental Table 5). We confirmed that *GNA15* expression was ∼60% reduced in the Gα15 knockdown group compared with the control group (Fig. 4B). Since Gα15 belongs to the Gαq/11 family and activates the downstream signaling pathways of Gαq/11 family, knocking down Gα15 should reduce the expression of Gαq-responsive genes. Several genes previously reported to be mediated by Gαq signaling—including *EGR1*, *DUSP5*, *LIF*, *ATF3*, and *SERPINE1*—were significantly downregulated in Gα15 knockdown group compared with control groups under clenbuterol-treated conditions^75^ (Fig. 4B), supporting the disruption of the Gαq signaling pathway by Gα15 knockdown in the RNA-seq.

Next, we identified genes that are mediated by the β2AR-Gαs and Gα15 signaling pathways, especially Gα15, which is understudied in β2AR signaling. The DEA demonstrated that knockdown of Gαs and Gα15 elicited opposing regulatory effects on a subset of genes, whereas another subset displayed similar regulatory responses. (Fig. 4C). To identify β2AR-Gαs and Gα15 mediated genes, we focused on genes with opposite expression patterns. Those upregulated by clenbuterol stimulation but downregulated upon loss of Gαs or Gα15, and conversely, those downregulated by clenbuterol but upregulated in the knockdown conditions. Notably, A distinct subset of genes exhibited concordant regulation by both Gαs and Gα15 signaling pathways (Fig. 4D). In contrast, a substantial number of genes were exclusively responsive to Gαs signaling (Fig. 4E), whereas only a small number of genes appeared to be uniquely mediated by Gα15 signaling. For instance, *CDCP1*, *CCL20*, *MAP2K3*, *DGKA*, *HS3ST3B1*, *NT5E*, and *MINK1* were upregulated following clenbuterol stimulation but downregulated upon Gα15 knockdown. Conversely, the expression of several genes, including *GCNT1*, *XK*, *LRRC4*, *TGFBR1*, *RRAGC*, *DAB2*, *GNAQ*, and *CUEDC1*, were reduced in response to clenbuterol treatment, yet were upregulated upon loss of Gα15 (Fig. 4F). These genes might be uniquely mediated by β2AR-Gα15 signaling in THP-1-derived macrophage-like cells. Although the magnitude of differential expression for these genes is modest and raises questions about biological relevance, one plausible explanation is ligand concentration-dependent coupling bias. At the 1μM clenbuterol concentration, β2AR signaling is likely dominated by Gαs engagement, which could mask or dilute the transcriptional profile driven by Gα15 and thereby result in loss or smaller fold changes for Gα15-mediated genes. Interestingly, *IL10* and *IL1b* expression was not reduced in either the Gαs or Gα15 knockdown groups, whereas both cytokines were downregulated in the GαsBP2 group. This finding suggests that the Gαs-cAMP signaling pathway may be associated with the regulation of *IL10* and *IL1β* expression, and compensatory signaling mechanisms may engage in chronic depletion of Gαs that preserve *IL10* and *IL1b* expression. Taken together, these exploratory findings suggest that Gαs and Gα15 are associated with partially overlapping but distinct subsets of β2AR-responsive genes. The Gα15-associated gene subset may reflect a non-redundant signaling contribution of Gα15 to β2AR-mediated macrophage-related transcriptional responses.

To further characterize the downstream effects of Gαs and Gα15 signaling, we performed GSEA for Gαs and Gα15 knockdown under clenbuterol stimulation with hallmarks from MSigDB, and list top ten hallmarks that is either positive or negative enriched in loss of Gαs and Gα15 based on the enrichment score (Fig. 5A & B). Loss of Gαs markedly enriched hallmarks associated with inflammatory responses, including the interferon-α and interferon-γ hallmark, while concurrently negatively enriched in several hallmarks typically linked to non-inflammatory activity or tissue homeostasis, such as epithelial-mesenchymal transition and hypoxia^76, 77, 78^. Loss of Gα15 showed several overlapping Hallmark enrichment patterns with loss of Gαs, while also exhibiting distinct enrichment differences. In GSEA, loss of Gα15 was associated with negative enrichment of MYC target and positive enrichment of apoptosis, oxidative phosphorylation, and protein secretion hallmark. By contrast, loss of Gαs was associated with positive enrichment of MYC target and KRAS signaling Hallmark gene sets. These results suggest that Gαs and Gα15 are associated with partially overlapping Hallmark enrichment patterns under β2AR stimulation, while also contributing to distinct transcriptional outcomes.

**Figure 5.**
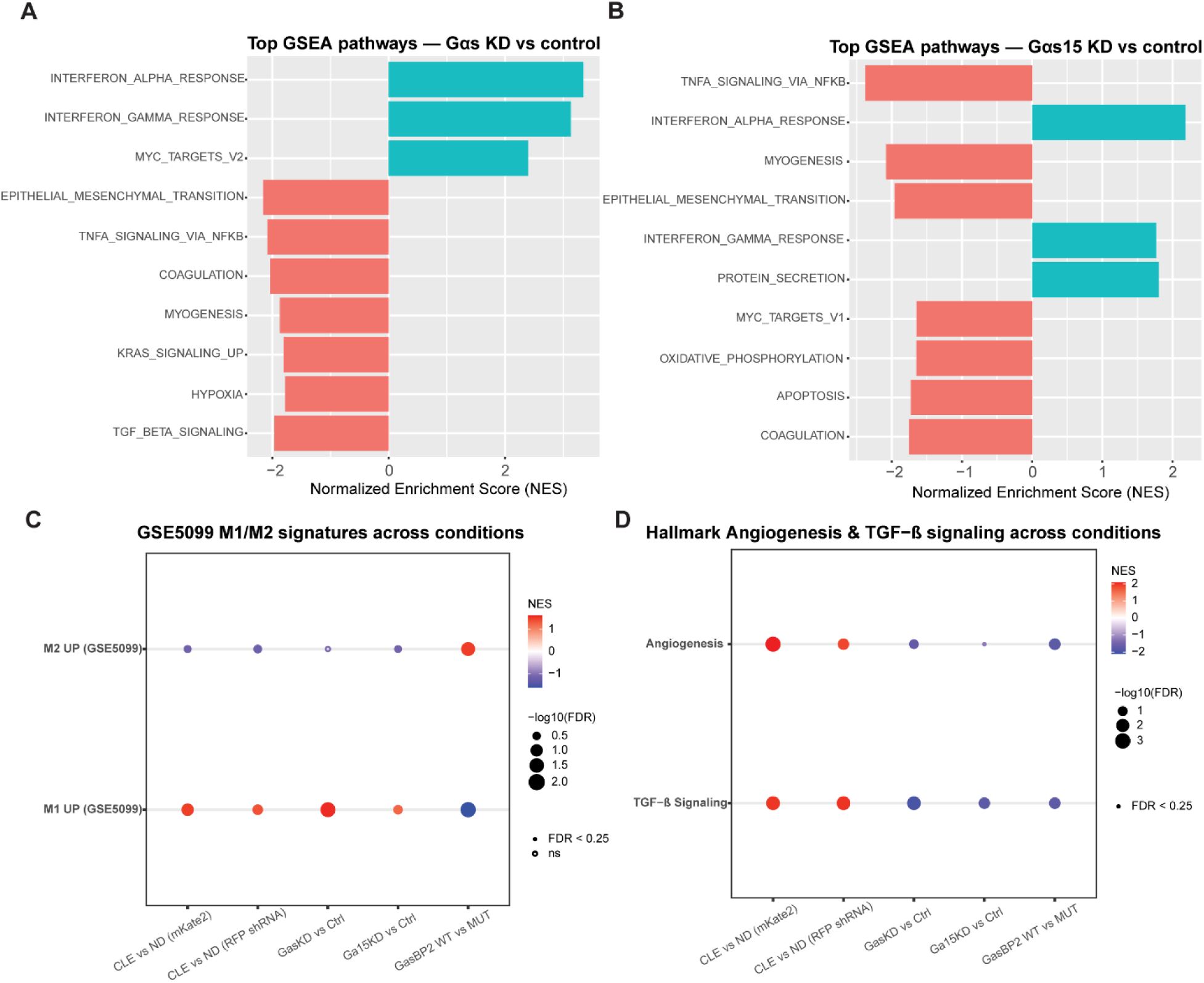
GSEA of M1/M2, TGF-β, and angiogenesis gene sets in Gαs- and Gα15-knockdown cells under clenbuterol treatment. (A) GSEA on the Gαs knockdown group versus control under clenbuterol stimulation. (B) GSEA on the Gα15 knockdown group versus control under clenbuterol stimulation. (C) GSEA on clenbuterol treatment versus vehicle, as well as the Gαs-, Gα15-knockdown, and GαsBP2 WT group versus control under clenbuterol stimulation, using the predefined M1 vs M2 gene sets (GSE5099) from MSigDB. (D) GSEA on clenbuterol treatment versus vehicle, as well as the Gαs-, Gα15-knockdown, and GαsBP2 WT group versus control under clenbuterol stimulation, using the predefined the hallmark of TGF-β signaling and angiogenesis from MSigDB. FDR values < 0.25 were considered significant.

M2 macrophages have been demonstrated to improve cardiac repair and preserve heart function after MI^79^. Furthermore, genetic ablation of β2AR specifically in immune cells has been reported to reduce survival following myocardial infarction and to diminish immune-cell infiltration to the injured myocardium^41^. To explore whether β2AR signaling as well as Gαs and Gα15 signaling pathways are associated with macrophage transcriptional profile relevant to M1 or M2 polarization and tissue repair, we performed GSEA for clenbuterol treatment, Gαs and Gα15 knockdown, and GαsBP2 inhibition. Gene sets were selected from predefined MSigDB collections, including immunologic signatures and Hallmark. We particularly examined macrophage M1/M2 gene sets from GSE5099^80^, as well as angiogenesis and TGF-β signaling hallmarks, which contain genes previously associated with tissue repair-related processes. Here, M1- and M2-associated signatures refer to predefined gene sets derived from classically activated pro-inflammatory macrophages and alternatively activated anti-inflammatory or tissue-reparative macrophages, respectively. In Figure 5, M1_UP and M2_UP denote macrophage M1 and M2 gene sets from GSE5099. Thus, enrichment of these signatures indicates transcriptional similarity to M1 or M2 macrophage, rather than direct evidence of functional macrophage polarization. Clenbuterol treatment was associated with positive enrichment of the M1_UP across control backgrounds and modest negative enrichment of the M2_UP gene set, suggesting transcriptional similarity to an M1 macrophage. Gαs knockdown was also associated with positive enrichment of M1_UP genes and negative, nonsignificant enrichment of M2_UP genes under clenbuterol-treated conditions. In contrast, blockade of Gαs-cAMP axis by GαsBP2 was associated with negative enrichment of M1_UP and positive enrichment of M2_UP. Gα15 knockdown was associated with positive enrichment of M1_UP and negative enrichment of M2_UP gene set (Fig. 5C). In addition, clenbuterol treatment was associated with positive enrichment of the TGF-β signaling and angiogenesis hallmark. These enrichments were reversed after Gαs or Gα15 knockdown and after GαsBP2 expression, with only a modest reduction in angiogenesis enrichment observed in the Gα15 knockdown condition (Fig. 5D). Together, these exploratory GSEA results suggest that β2AR agonism, Gαs and Gα15 perturbation, and GαsBP2 expression are associated with distinct macrophage M1 and M2 polarization- and tissue repair-related gene sets in THP-1-derived macrophage-like cells.

## 4. Discussion

β2AR canonically couples to Gαs and had been demonstrated to couple to Gα15 with epinephrine. However, whether other β2 agonists also perform the couple of Gα15 remains uncharacterized. Here, we tested four other agonists on Gα15 activation and compared with Gαs. We observed that four agonists activate Gα15 with consistently higher potency than Gαs in TRUPATH triple assays. This potency bias suggests that β2AR may facilitate GDP-GTP exchange of Gα15 more efficiently than Gαs. Gα15 may act as more sensitive transducer for β2AR in immune cells. In addition, terbutaline may be a partial agonist for Gαs, but a full agonist for Gα15, compared with epinephrine. In addition to β2AR agonists, we tested three clinically used β-blockers to investigate how β-blocker block β2AR-mediated Gα15 activation. Our findings suggest that timolol may be a more potent β-blocker for both Gαs and Gα15 compared with carvedilol and labetalol. Moreover, timolol may differ in its blockade on β2AR-mediated G protein activation in TRUPATH triple assay. One possible explanation for these observations could be that different agonists and β-blockers may bind β2AR with distinct affinities or stabilize different receptor conformations, which could influence coupling to individual G proteins and contribute to differences in apparent potency or efficacy^81^. However, since these observations were obtained using a single assay platform and limited sample size, they should be interpreted cautiously and considered as exploratory data. Although identical transfection conditions were used, biosensor expression was not independently quantified, potency and efficacy differences derived from TRUPATH triple assay may be influenced by biosensor expression levels. Orthogonal assays will be required to determine whether these effects reflect true ligand-specific modulation of β2AR-G protein coupling. Besides, most reported beneficial effects of β-adrenergic ligands have been attributed to canonical Gα-cAMP signaling, whereas the potential contribution of Gα15 remains unclear. Our findings raise the possibility that β2AR-Gα15 signaling may also affect the impact of β2AR drugs tested in this paper in immune system, but this signaling pathway has not been linked to clinical benefit.

Clenbuterol-induced transcriptional profile was associated with positive enrichment of predefined hallmarks, including inflammatory response, TNF signaling and IL-2/STAT5 signaling, as well as tissue repair-associated hallmark, such as TGF-β signaling and angiogenesis. GO enrichment analysis also identified terms associated with chemotaxis and small GTPase-mediated signal transduction, suggesting that β2AR activation may be linked to transcriptional programs involved in cell migration, cytoskeletal remodeling, or phagocytic activity. This is consistent with previous reports showing that β2AR deficiency can impair immune cell recruitment^41, 82^ and stimulation of β2AR enhance phagocytosis^83^. Transcriptome analysis following Gαs and Gα15 knockdown suggested that these pathways are associated with partially overlapping but distinct transcriptional responses in macrophage-like cells. Gαs knockdown produced broader transcriptional changes than GαsBP2 expression. This difference may reflect broader effects of reducing total Gαs abundance compared with selectively perturbing the canonical Gαs-cAMP axis by GαsBP2. One possible explanation is that Gαs may influence transcription response through cAMP-independent signaling pathways in addition to the canonical Gαs-cAMP pathway. However, since cAMP production was not directly measured in the RNA-seq samples, these data should not be interpreted as a direct comparison of Gαs knockdown versus GαsBP2 inhibitory efficacy.

GSEA showed that clenbuterol treatment was associated with an M1 transcriptional profile together with enrichment of TGF-β signaling and angiogenesis hallmark. Gαs knockdown was associated with strong positive enrichment of the M1 profile, whereas Gα15 knockdown was associated with positive M1 enrichment and negative M2 enrichment. In contrast, GαsBP2 expression showed an opposing enrichment pattern. These findings suggest that β2AR-associated transcriptional regulation may reflect integration of multiple G-protein-dependent inputs, including Gαs-cAMP- and Gα15-dependent pathways, resulting in the net transcriptional profile observed after β2AR activation. However, since these analyses are based on curated gene set enrichment, they should be interpreted as transcriptional associations rather than direct functional evidence of immune responses, M1/M2 polarization, or tissue repair. The GαsBP2-associated M1/M2 enrichment pattern differs from previous studies linking cAMP signaling to M2-like macrophage polarization^84, 85, 86^. This discrepancy may reflect differences in cell model, stimulation context, timing, or transcriptome-wide analysis versus selected markers. Particularly, our experiments used clenbuterol stimulation in THP-1-derived macrophage-like cells without additional polarizing cytokines, such as IL4 or LPS/IFNγ. Additional limitations include the use of THP-1-derived macrophage-like cells rather than primary macrophages and potential compensatory effects for chronic Gαs or Gα15 knockdown. Future studies directly measuring macrophage cytokine secretion, chemotaxis, phagocytosis, and tissue-repair-related activity will be required to determine the functional significance of these transcriptional responses. Furthermore, β2AR also couples to Gαi/o and recruits β-arrestins in macrophages, which is not being investigated in this study. Several studies have demonstrated that β2AR can regulate immune responses through noncanonical pathways, including ERK1/2, p38, and MAPK signaling via Gαi/o and β-arrestins^87, 88, 89^. However, whether these signaling pathways promote pro- or anti-inflammatory activity of macrophages remains incompletely defined and is likely context- and time-dependent. Thus, future study should also include these two signaling pathway to obtain a more comprehensive understanding of β2AR signaling in macrophages. In addition to β2AR, β3AR abundance has been reported to increase in failing heart^90^. Studies have demonstrated that β3AR activation can induce nitric oxide signaling pathways in both cardiomyocytes and endothelial cells to perform cardioprotection^91, 92^; however, β3AR signaling has been demonstrated mainly in cardiomyocytes. Nevertheless, whether β3AR signaling in macrophages contributes to cardiac protection and repair remains unknown and needs to be further investigated.

In summary, this study explored a previously underappreciated β2AR signaling pathway involving Gα15 in macrophage-like cells. Our in vitro pharmacological profiling suggests that β2AR agonists and β-blockers can display distinct apparent activity profiles across Gαs and Gα15. Transcriptomic analyses further suggest that Gα15 may contribute to M2-associated transcriptional profile under β2AR stimulation, whereas the Gαs-cAMP signaling axis showed an opposing M1/M2-associated transcriptional pattern. Overall, these findings highlight potential differences between canonical β2AR-Gαs signaling and noncanonical β2AR-Gα15 signaling, providing a foundation for future studies investigating the role of β2AR-Gα15 signaling in macrophage function.

## Abbreviation

β2AR (β2-adrenergic receptor), BRET (bioluminescence resonance energy transfer), PMA (phorbol 12-myristate 13-acetate), DEA (differential expression analysis), DEG (differentially expressed gene), GSEA (gene set enrichment analysis), GO (gene ontology) RFP (red fluorescent protein), NES (normalized enrichment score), MSigDB (molecular signature database)

## Acknowledgments

We thank Dr. Roh-Johnson (The University of Utah) for the THP-1 cell line and the lentiviral plasmids and Dr. Garcia-Marcos (Boston University) for the GαsBP2 wild-type and mutant plasmids. Figure 3A and supplemental figure 3A were made using BioRender.

## Data availability

The authors declare that all the data supporting the findings of this study are contained within the paper. The RNA-seq data are available in NCBI Gene Expression Omnibus (GEO) with reference number GSE325719. All other data will be made available from the corresponding author upon reasonable request.

## CRediT authorship contribution statement

**Yuan-En Sun**: Conceptualization, Formal analysis, Investigation, Writing – original draft. **Qing Li**: Data curation, Formal analysis. **Justin G. English**: Conceptualization, Funding acquisition, Resources, Supervision, Writing – review and editing.

---

**Supplemental Figure 1.**
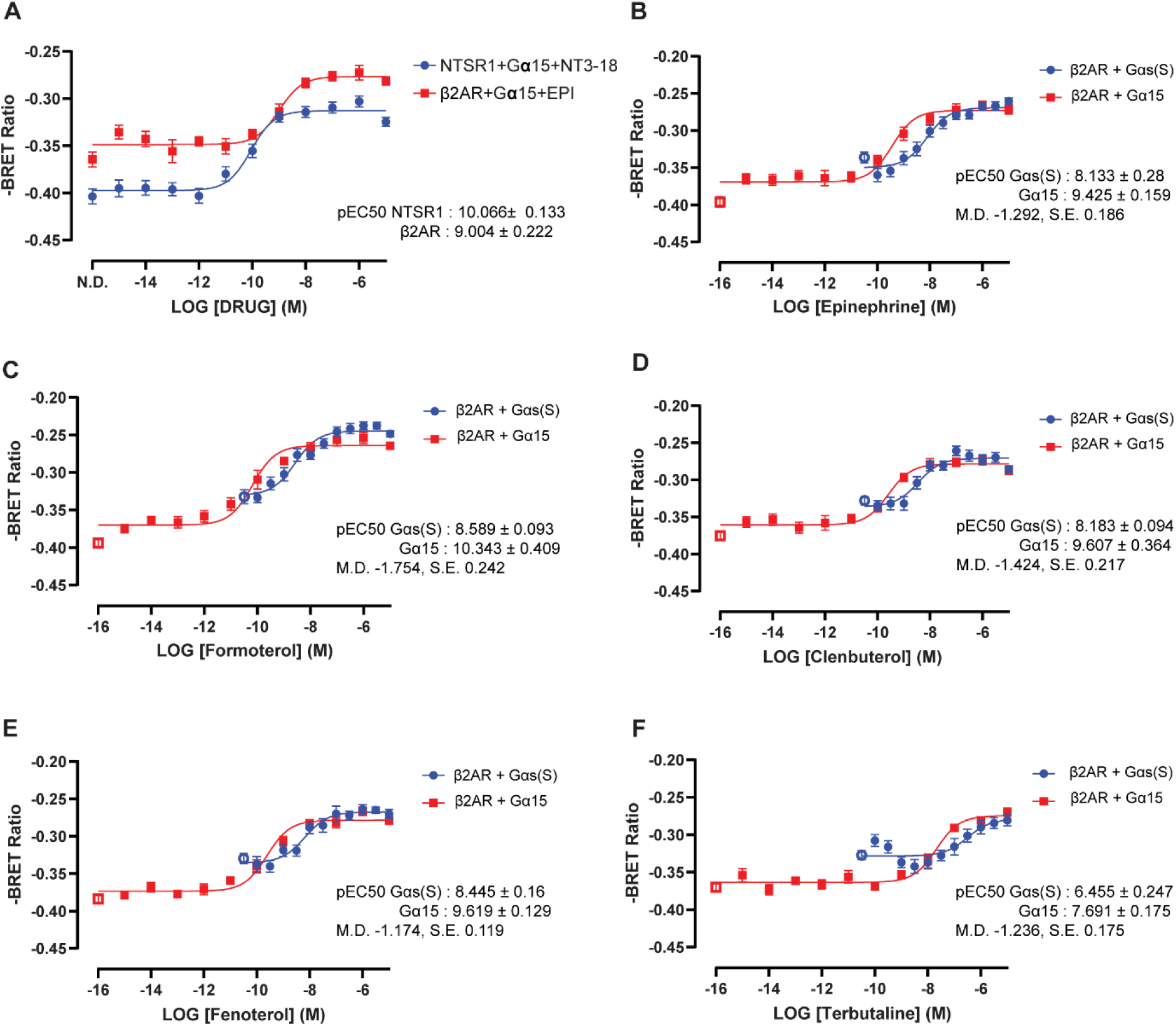
TRUPATH concentration-response analysis of Gαs and Gα15 signaling via β2AR and NTSR1. (A) Raw BRET ratio of Gα15 TRUPATH concentration-response curves of NTSR1 and β2AR with endogenous agonist, neurotensin 8-13 and epinephrine, respectively. Data are represented as mean values ± SEM, N=3, n=4. (B) Raw BRET ratio of Gαs and Gα15 TRUPATH concentration-response curves of β2AR with epinephrine. (C) Formeterol (D) Fenoterol (E) Clenbuterol (F) Terbutaline. Hollow dot represents no drug (N.D.). Data are represented as mean values ± SEM, N=3, n=4. N refers to biological replicate, and n refers to technical replicate. pEC50 values are presented as mean ± SD from three independent biological replicates.

**Supplemental Figure 2.**
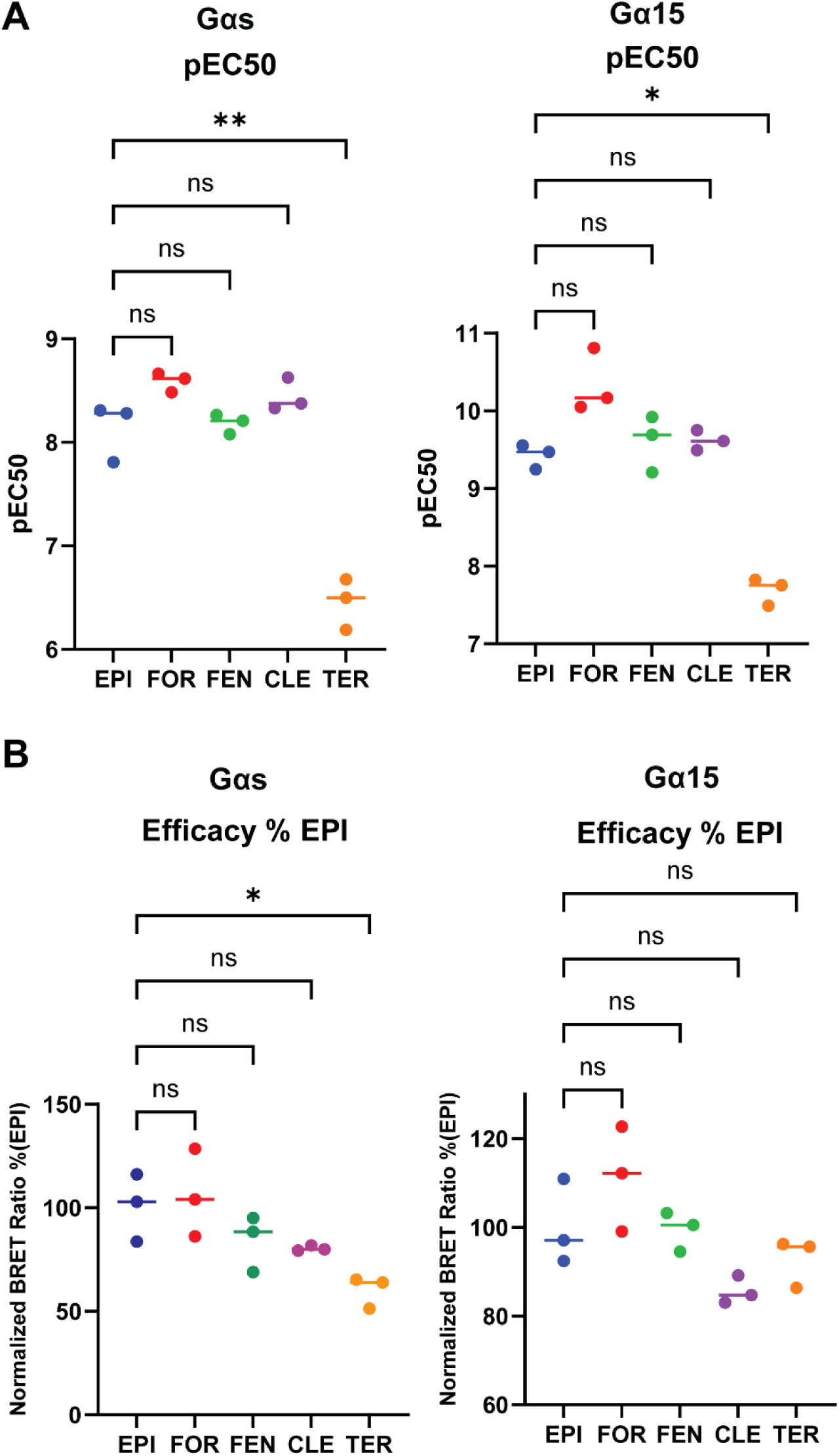
Statistical analysis of pEC50 and Efficacy of tested β2AR agonists versus epinephrine. (A) Statistical analysis of pEC_50_ (B) Statistical analysis of efficacy. Data are represented as mean ± SD, N=3. One-way ANOVA followed by Dunnett’s test, P-value: 0.1234(ns), 0.0332(*), 0.0021(**), 0.0002(***) & 0.0001(****). N refers to biological replicate.

**Supplemental Figure 3.**
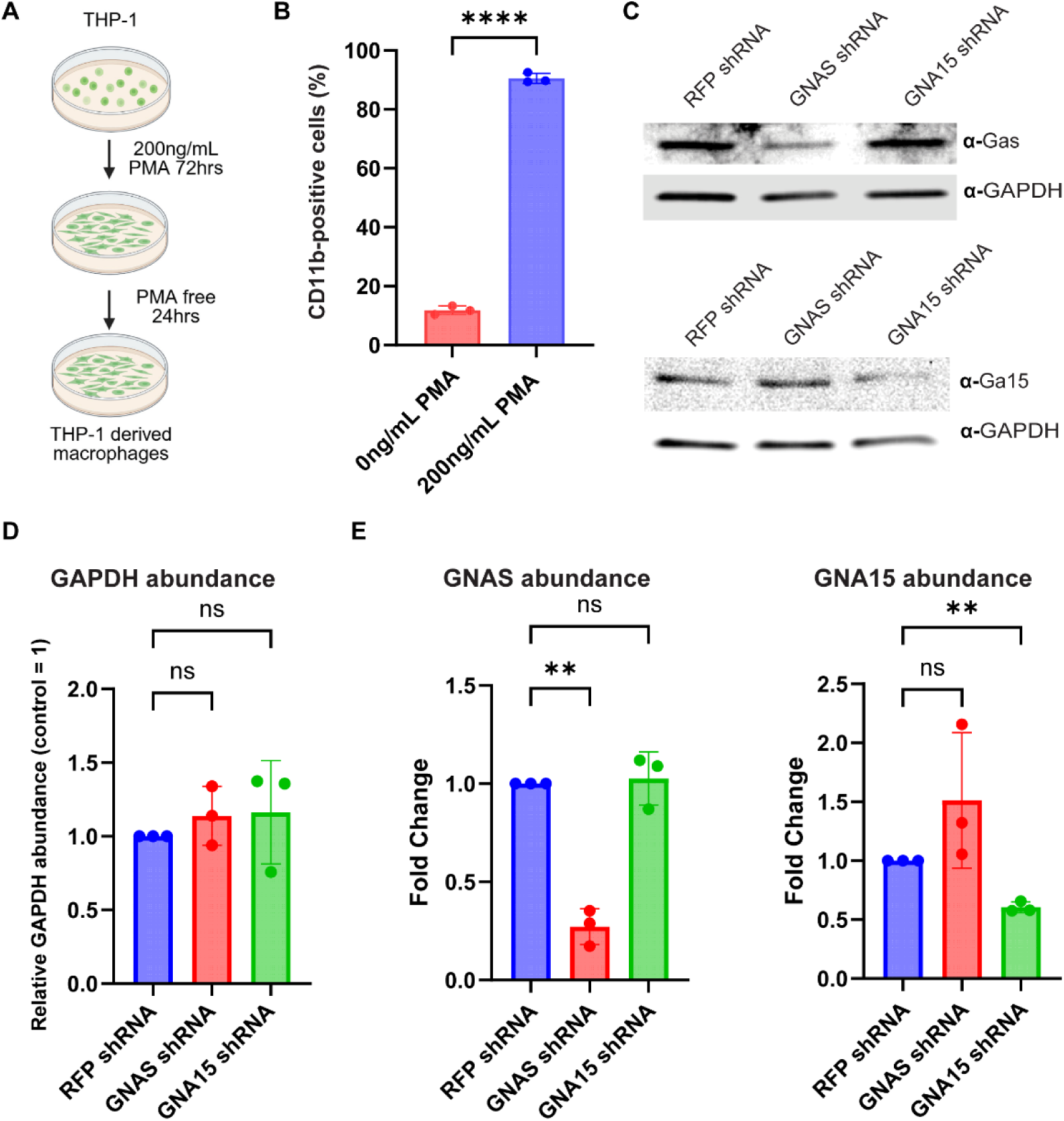
Knockdown of Gαs and Gα15 in THP-1-derived macrophage-like cells. (A) Workflow used to differentiate THP-1 cells into macrophage-like cells using PMA. (B) Quantification of CD11b-positive cells. N=3, t-test. (C) Upper: representative immunoblot of Gαs knockdown in cells with shRNA by lentiviral transduction. Bottom: representative immunoblot of Gα15 knockdown in cells with shRNA by lentiviral transduction. (D) Quantification of GAPDH abundance in cells after shRNA transduction 4 days. (E) Quantification of Gαs and Gα15 abundance in cells after shRNA transduction for 4 days. Data represents as mean ± SD, N=3, One-way ANOVA by Dunnett’s test, P-value: 0.1234(ns), 0.0332(*), 0.0021(**), 0.0002(***) & 0.0001(****). N refers to biological replicate.

**Supplemental Figure 4.**
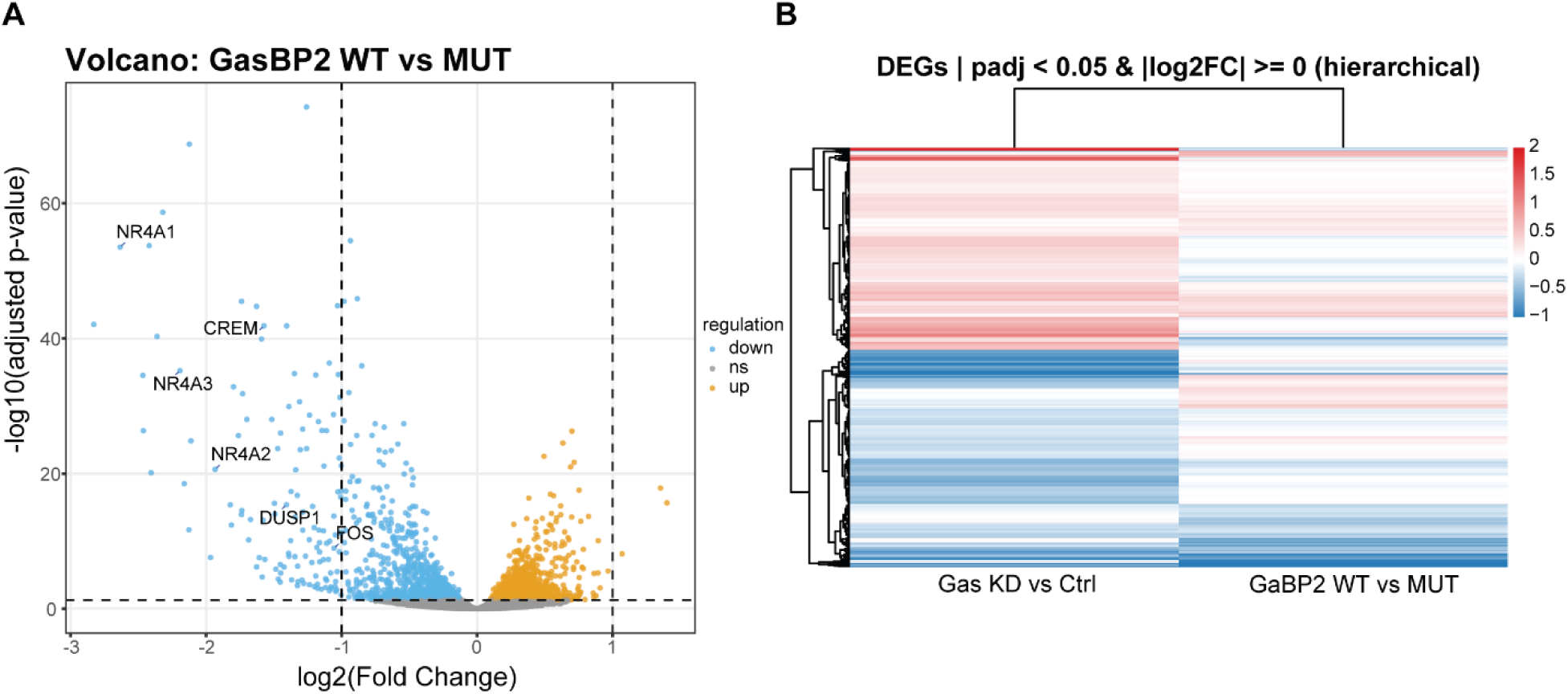
Transcriptomic Profiling of Gαs-cAMP-dependent signaling in response to clenbuterol in macrophage-like cells. (A) Volcano plot of DEGs between the GαsBP2 WT group versus GαsBP2 mutant control under clenbuterol treatment, with genes known to be regulated by the β2AR-Gαs-cAMP-CREB pathway annotated, defined by ANOVA with an adjusted p-value less than 0.05 (yellow and blue dots). (B) Hierarchical clustering heatmap of DEGs between Gαs knockdown versus control and GαsBP2 wild-type versus mutant control, defined by ANOVA with an adjusted p-value < 0.05.

